# The Pivotal Involvement of the *Respiratory burst oxidase G* (*SlRbohG*) Gene in H_2_O_2_ Production Under Stress for Proper Na^+^ Homeostasis Regulation in Tomato

**DOI:** 10.1101/2024.03.12.584686

**Authors:** I Egea, T Barragán-Lozano, Y Estrada, M Jáquez-Gutiérrez, FA Plasencia, A Atarés, B Garcia-Sogo, C Capel, F Yuste-Lisbona, JM Egea-Sánchez, FB Flores, T Angosto, M Moreno, R Lozano, B Pineda

**Affiliations:** Centro de Edafología y Biología Aplicada del Segura, Consejo Superior de Investigaciones Científicas, 30100 Espinardo, Murcia, Spain; Instituto de Biología Molecular y Celular de Plantas (UPV-CSIC), Universidad Politécnica de Valencia, 46011 Valencia, Spain; Centro de Investigación en Biotecnología Agroalimentaria (CIAIMBITAL), Universidad de Almería, 04120 Almería, Spain

**Keywords:** tomato, *Solanum lycopersicum*, *SlRbohG* gene, NADPH-oxidase, H_2_O_2_ production, salt stress, Na^+^ homeostasis, aquoporins, water flux

## Abstract

Regulation of sodium homeostasis is crucial for plant response to salinity conditions. Here we report on the genetic and physiological characterization of two tomato allelic mutants, *<u>s</u>odium <u>ga</u>therer1-2* (*sga1-2*), which exhibit pronounced chlorosis and hyperhydration under salt stress. Mapping-by-sequencing revealed that mutant phenotype resulted from mutations in the *SlRbohG* gene, and CRISPR/Cas9 knockouts of this gene gave phenocopies of the *sga1-2* mutants. Physiological analyses showed that *sga1-2* salt hypersensitivity is linked to an increase of Na^+^ and water transport from roots to shoots, which explains their extreme chlorosis and hyperhydration under salinity conditions. At the molecular level, *SlPIP2;12* gene, an aquaporin down-regulated in the WT under salt stress, was overexpressed in the *sga1-2* mutants, which could enhance water transport to the shoot. Also, *sga1-2* mutants exhibited a significant reduction in the expression of key sodium transporters, thus modifying the normal distribution of Na^+^ in tomato plant tissues. Furthermore, treatment of WT plants with the NADPH oxidase inhibitor DPI prevented H_2_O_2_ production in response to salinity, resulting in elevated Na^+^ accumulation in the shoot and reduced expression of the *SlHKT1;2* gene in root. Altogether, our results show that *SlRbohG* plays a central role in salt tolerance through ROS-mediated signaling.

**HIGHLIGHT:** Loss of function of tomato *SlRbohG* gene leads hypersensibility to salt stress due to increased Na^+^ and water transport from root to shoot.

## INTRODUCTION

Great efforts are currently being invested in increasing abiotic stress tolerance in species of agronomic interest to cope with the imminent effects of climate change and global warming. Among abiotic stresses, salinity is one of the most limiting to crop productivity, and the area affected by salt stress continues to grow (**Munns *et al*., 2020; Corwin, 2021**). However, generating tolerant crops remains a challenge due to the complexity of the trait and, above all, the lack of knowledge about the key genes and the molecular and physiological processes that determine tolerance (**Bowerman *et al*., 2023**). One of the main bottlenecks in breeding programs is the strong reduction of phenotypic and genetic diversity in modern germplasms, due to domestication itself firstly and the development of commercial cultivars secondly, which has led to the phenomenon of genetic erosion, as recently remarked by **Kromdijk and McCormick (2022)**. In fact, the selection of genes related to agronomic traits during the domestication process (**Liang *et al*., 2021**) was accompanied by the loss of alleles involved in abiotic stress tolerance (**Zhang *et al*., 2023**). Therefore, the identification and introduction of genetic determinants that were lost during domestication may facilitate the development of salt-tolerant crops (**Eckardt *et al*., 2023**). Tomato (*Solanum lycopersicum* L.) is not only an essential crop for its economic importance and high nutritional value, but it is also a model dicot for advancing the understanding of the underlying mechanisms of abiotic stress tolerance (**Seymour and Rose, 2022)**. Salinity causes damage to plants by osmotic stress (due to reduced water availability), ionic stress (mainly as a consequence of the toxic effect of Na^+^) and accumulation of reactive oxygen species (ROS). In tomato, salt tolerance is greatly determined by its ability to regulate Na^+^ transport from root to shoot over time, and to store it in a non-toxic manner into the vacuoles of root and leaf cells (**Egea *et al*., 2018**). The regulation of Na^+^ homeostasis in tomato under salt stress conditions is mediated by the coordination of several Na^+^ transporter, being tomato a very good plant model to study long distance transport of saline ions because of its physiology and anatomy (**Egea *et al*., 2022**).

In addition, the generation of ROS is a crucial process in the response of plants to abiotic stress. Oxidative stress typically comes as a secondary component after primary stresses such as salinity. In recent years, the double role of ROS has been well-established, as ROS are considered harmful oxidants at high concentrations, while transient and compartment-specific ROS accumulation serves as signals for various plant developmental processes as well as responses to stress (**Castro *et al*., 2021**). In the apoplast, ROS are produced through the activation of ROS-producing enzymes such as apoplastic peroxidases, polyamine oxidases, and plasma membrane-localized NADPH oxidases (respiratory burst oxidase homologs, Rbohs). NADPH oxidases are considered the most important class of apoplastic ROS-producing enzymes, generating superoxide (O_2_^.-^) from molecular oxygen (O_2_). Evidence shows that ROS plays an essential role in plant salinity tolerance (**Hossain and Dietz, 2016; Niu *et al*., 2018**). For example, Arabidopsis mutants with *Rboh* loss-of-function alleles have been shown to be hypersensitive to salt stress, supporting the critical role of these genes in the salt stress response. Thus, **Ma *et al*. (2012)** observed that the Arabidopsis *AtrbohD1/F1* and *AtrbohD2/F2* double mutants displayed significantly inhibited generation of ROS and, as a consequence, exhibited higher Na^+^/K^+^ ratios than WT. Similarly, **Jiang *et al*. (2013)** reported that Arabidopsis *AtrbohF* knockout mutants showed an increased salt sensitivity and impaired Na^+^/K^+^ homeostasis. Furthermore, a recent report showed that expression patterns of the *RBOHD* and *RBOHF* genes under salinity differ between the glycophyte Arabidopsis and the halophyte *Eutrema salsugineum* (**Pilarska *et al*., 2021**). In tomato, phylogenetic analyses with the Arabidopsis and rice *Rbohs* genes indicated that the tomato *SlRbohs* genes are classified into five subgroups (**Li *et al*., 2015**). *SlRbohG*, previously reported as *SlRboh1* (**Sagi *et al*., 2004**), belongs to subgroup IV, where Arabidopsis RbohF and rice OsRbohA/C homologues are included. *SlRbohG* gene is involved in tolerance to salt stress. Thus, **Yi *et al*. (2015**) studied the impact of high atmospheric CO_2_ concentrations on the responses of tomato plants to salt stress, with a particular focus on the role of *SlRbohG*. These authors demonstrate that high CO_2_ can counteract the negative impact of salt stress on photosynthesis and biomass production in an apoplastic H_2_O_2_-dependent manner. Furthermore, the regulation of H_2_O_2_ also contributed to the regulation of Na^+^ transport from roots to shoots, a process influenced by stomatal movement and Na^+^ delivery from xylem to leaf cells. However, more advances are necessary to elucidate the roles of *SlRboh* genes in the salt tolerance of this crop species (**Raziq *et al*., 2022**).

In this research work, we report the identification of two salinity hypersensitive allelic mutants, named *<u>s</u>odium <u>ga</u>therer1* and *2* (*sga1-2*), whose most singular traits under salt stress conditions are leaf chlorosis and hyperhydration due to Na^+^ and water over-accumulation in shoot. By combining mapping-by-sequencing and CRISPR/Cas9 genome editing methods, we revealed that the salt hypersensitivity phenotype of the mutants is due to the loss of function of the *SlRbohG* gene. Here, we show the first results unraveling the crucial role of *SlRbohG* in long-term salt tolerance of tomato. The physiological and molecular characterization of the two mutants indicates that the primary role of *SlRbohG* in salt conditions is the regulation of Na^+^ transport from root to shoot mediated by the H_2_O_2_ production.

## MATERIAL AND METHODS

### Screening and identification of *sga1* and *sga2* tomato mutants

The *sga1* and *sga2* mutants were identified in a collection of T-DNA mutants of tomato cv Moneymaker (**Pérez-Martín *et al*., 2017**) during the *in vitro* screening of T1 segregating families (30-40 plants per family) grown in basal culture medium supplemented with 100 mM NaCl (**Egea *et al*., 2018**). Additional experiments in the same conditions but with different NaCl concentrations allowed to confirm the phenotypical reproducibility of both mutants. In order to estimate the number of inserts bearing a functional *nptII* marker gene, a segregation analysis in kanamycin-containing medium consisting of Murashige and Skoog (MS) salts (**Murashige and Skoog, 1962**), sucrose (10 g L^-1^), and kanamycin (100 mg L^-1^) was carried out. A co-segregation analysis was performed as described by **Jáquez-Gutiérrez *et al*. (2019)** to study the association between the mutant phenotype and the *nptII* gene expression. For this purpose, after identification of the mutant in NaCl-supplemented medium, shoot-apex from both the WT and mutant seedlings were cut off and transferred to MB3 medium (**Moreno *et al*., 1984**) supplemented with kanamycin (50 mg L^-1^). In this medium, the non-expressing *nptII* shoots do not develop either roots or true leaves, while those expressing *nptII* develop adventitious roots and true leaves. Finally, to determine whether the two recessive mutations were alleles of the same gene or different genes, a complementation test was carried out by crossing the two mutants and evaluating the phenotype of the progeny under *in vitro* conditions in a medium supplemented with NaCl.

### Molecular cloning of the *sodgat* mutations

A mapping-by-sequencing approach was used to identify the mutations underlying the *sga* locus, according to the procedure described by **Yuste-Lisbona *et al*. (2021).** Briefly, an F_2_ mapping population was generated by self-fertilizing a F_1_ plant resulted from a cross between a *sga2* mutant plant without T-DNA insertion and a *S. pimpinellifolium* LA1589 accession plant. Equal amounts of DNA from 30 randomly selected phenotypically wild-type and 25 phenotypically mutant plants were pooled to construct wild-type and mutant pools, respectively. DNA pools were sequenced with paired-end reads on Illumina HiSeq2000 platform (Illumina, Inc., San Diego, CA, USA) and the obtained sequences have been deposited at the Sequence Read Archive database at the National Center for Biotechnology Information under BioProject accession number PRJNA860761. Reads were aligned to the tomato genome reference sequence v.4.0 (ITAG4.0) using Bowtie2 with the default parameters (**Langmead and Salzberg, 2012**). The allele frequency ratio for biallelic variants was calculated as nonreference allele counts/total allele counts. The average allele frequencies were plotted along each chromosome using a sliding window and step size of 1000 and 100 variants, respectively. Variant analysis filtering criteria on the candidate genomic region to host the *sodgat2* mutation was based on (i) the alternative allele must be in a heterozygous (0/1) and homozygous (1/1) state in wild-type and mutant pools, respectively; and (ii) alternative alleles must not have been previously reported in the sequenced tomato genomes (**Lin *et al*., 2014; The 100 Tomato Genome Sequencing Consortium, 2014**). The translocation breakpoints identified to be underlying the *sga2* mutation were validated by PCR using the primers listed in **Supplementary Table S1**. Afterwards, a co-segregation analysis between the *sga2* mutant phenotype and the translocation breakpoints was performed in the F_2_ population using the Ch08-F and Ch08-R or Ch10-F and Ch10-R primers for translocation in chromosome 08 or 10, respectively (**Supplementary Table S1**). Finally, the *SlRbohG* locus was sequenced in the *sga1* mutant by Sanger technology to determine the molecular nature of this mutation. The sequences of primers used are presented in **Supplementary Table S1**).

### Generation of CRISPR/Cas9 lines

CRISPR/Cas9 mutagenesis was performed according to the protocol described by **Vazquez-Vilar *et al*. (2016).** Briefly, Breaking-Cas web software (**Oliveros *et al*., 2016**) was used to design two single-guide RNA target sequences. The sgRNA1 (GCATGAACGCCGGTGGACGT) and sgRNA2 (GCTTGCGAGCAGCCAGAGGC) were designed at the beginning and end of the coding region of the *SlRbohG* gene (*Solyc08g081690*), respectively. Genetic transformation experiments were developed as described by **Ellul *et al*. (2003),** using *Agrobacterium tumefaciens* LBA4404 strain. The ploidy levels of transgenic plants were assessed by flow cytometry following the protocol described by **Atarés *et al*. (2011)**. Diploid first-generation transgenic (T_0_) plants were genotyped with primers that cover the target recognition region of the sgRNAs (**Supplementary Table S1**). PCR products were purified for cloning into the pGEMT vector (Promega, Madison, WI, USA). A minimum of 10 clones per PCR product were sequenced to characterize the edited alleles in the transgenic plants.

### Salt stress assays and physiological analysis

Tomato WT (cv. Moneymaker), and T3 homozygous plants of *sga1-2* mutants were used in all experiments. *In vitro* and *in vivo* assays were performed as previously described (**Jáquez-Gutiérrez *et al*., 2019; Egea *et al*., 2018**). To determine the relative importance of root and shoot in the mutant phenotype, grafting assays were performed as described by **Santa-Cruz *et al*. (2002)**.

Measurements of water contents, leaf stomatal conductance (g_s_), transpiration rate (E) and leaf osmotic potential (ψ_s_) were performed as described in **Egea *et al*. (2018)**. The concentrations of Na^+^ and K^+^ were analysed by inductively coupled plasma optical emission spectroscopy (ICP-OES) (Ionomics Platform of CEBAS-CSIC, Murcia, Spain).

### Gene expression analysis by RT-qPCR

The spatial and temporal expression of *SlRbohG* (*Solyc08g081690*) was analysed in WT plants grown in control and salt stress conditions. The expression levels of aquaporins (AQPs) genes of plasmatic membrane (PIPs) and tonoplast (TIPs) were analysed in root and leaf of plants grown in control and after 2 days of salt treatment (DST). Moreover, genes involved in Na^+^ homeostasis (*SlSOS1, SlHKT1;1* and *SlHKT1;2*), K^+^ homeostasis (*HAK5* and *SKOR*) and in Na^+^ vacuolar compartmentation (*SlNHX3* and *SlNHX4*) were analysed in control and salt treated roots after 2 DST.

Material was previously frozen in liquid N_2_ and stored at −80°C by quantitative RT-qPCR. All primers used for RT-qPCR are listed in **Supplementary Table S2**. Total RNA was isolated using a RNeasy kit (Qiagen) and RT-qPCR was performed as previously described (**Egea *et al*., 2018**). Relative expression data were calculated as described by **Asins *et al*. (2013)** using the tomato elongation factor 1α (LeEF1α, acc. AB061263) as housekeeping gene.

### Analysis of H_2_O_2_

Histochemical staining of H_2_O_2_ was assayed with 2,7-dichlorodihydrofluorescein diacetate (H_2_DCF; Invitrogen) as described by (**Potocký *et al*., 2007**). Briefly, 5 mm diameter leaf discs or 1 mm thick steam sections were incubated under darkness and vacuum conditions with a solution 60 μM of H_2_DCF in phosphate buffer 200 mM (pH 7,4) for 30 min. The tissues were imaged with confocal microscopy Leica DM6B motorized in X, Y, Z equipped with a Hamamatsu ORCA-flash 4.0LT fluorescence camera. Excitation 488 nm and emission 500–550 nm values were used. The same image parameters were manually adjusted to each tissue, to obtain a comparable fluorescence intensity between different treatments.

For the determination of H_2_O_2_ content by spectrophotometry, a modified ferrous ammonium sulphate/xylenol orange (FOX) method described by **Cheeseman (2006)** was used. Briefly, 300-500 mg of sample were dissolved in acidified acetone (25 mM H_2_SO_4_) buffer and mixed with the test eFOX solution (100 μM Xylenol orange, 250 μM SFeSO_4_, 100 μM Sorbitol and 1% ethanol) at least 30 minutes after measuring the difference in absorbance between 550 and 800 nm at RT using a using a plate reader PowerWave XS2 (BioTek). For each assay, the H_2_O_2_ content was quantified referring to an internal standard of a different dilution of H_2_O_2_ (30 % v/v).

### DPI (NADPH oxidase inhibitor) application assay

Diphenylene iodonium (DPI), an NADPH oxidase inhibitor, was used to analyse the potential role of the H_2_O_2_ production in regulating Na^+^ transport and its accumulation in roots and shoots (**Li *et al*., 2007**). A hydroponic assay was performed with WT seedlings. At the two-leaf stage, salt treatment (75 mM NaCl) was added to the nutrient solution. The salt treatment was applied alone, without DPI, in half of the seedlings, while DPI treatment (100 mM) was added to the other half of seedlings for 6 h. After this time period, irrigation solution was replaced by salt solution (75 mM NaCl) again and all seedlings were maintained until sampling. H_2_O_2_ accumulation was imaged by confocal microscopy in roots. The Na^+^ content was analysed in root and shoot after 2 days of salt treatment. Moreover, the expression of two main genes involved in Na^+^ homeostasis, *SlSOS1* and *SlHKT1;2*, was measured after 1 DST.

### Statistical analysis

Experimental data are presented as mean ± SE values. Statistical analysis was performed by Tukey’s test (p < 0.05) using the SPSS 24.0 software package. Significant differences between means are denoted by asterisks or letters.

## RESULTS

### Isolation and genetic characterization of the salt hypersensitive *sga* mutants

The *sodium gatherer 1-2 (sga1-2*) mutants were isolated from *in vitro* phenotypic screening of a collection of T1 segregating T-DNA lines (**Pérez-Martín *et al*., 2017**) performed under salt conditions (100 mM NaCl). Both mutants displayed a similar phenotype, characterized by abnormal symptoms of hyperhydration and intense chlorosis in different parts of the seedlings (**Fig. 1**). Interestingly, the hypersensitivity to salt stress of both mutants was observed *in vitro* at NaCl concentrations as low as 25 mM (**Supplementary Fig. S1).** Subsequently, the salt sensitivity of both mutants was corroborated *in vivo* by growing T1 plants under greenhouse conditions (**Supplementary Fig. S2**).

**Fig. 1.**
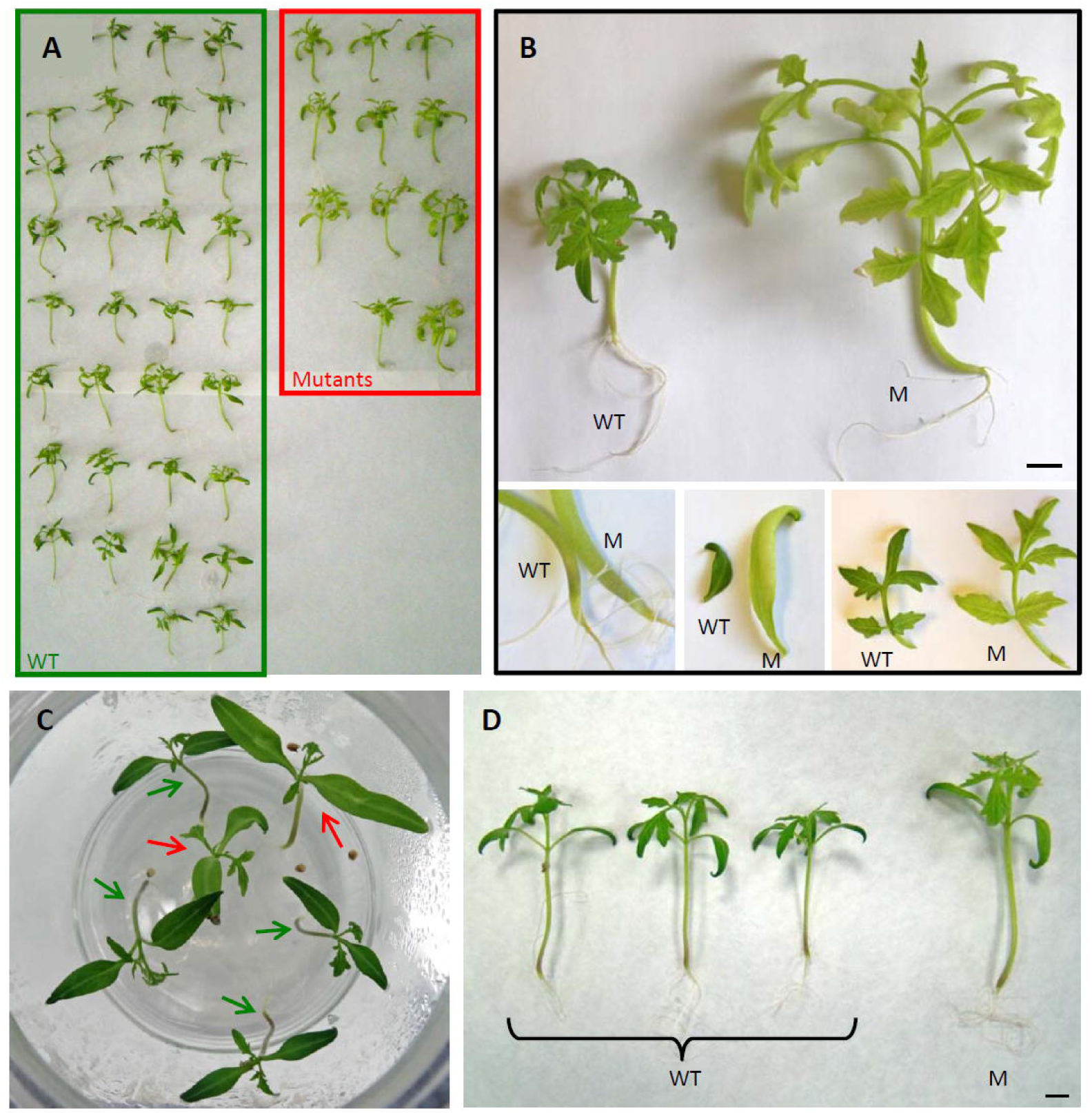
Identification of salt hypersensitive *sga1* and *sga2* mutants. A, Identification of the monogenic recessive *sdg1* mutant. B, Chlorosis and hyperhydration symptoms in *sdg1* mutant (bigger mutant size is related to hyperhydration symptoms), and close-up of hyperhydration in hypocotyl (bottom left), and chlorosis and hyperhydration in both cotyledon (bottom centre) and leaf (bottom right). C, Identification of the monogenic recessive *sga2* mutant. D, Chlorosis and hyperhydration symptoms in *sga2* mutant. Green and red arrows show WT and mutant plants, respectively. Plants of WT and *sga1* and *sga2* (C-D) mutants (M) were grown under salt stress conditions in vitro (100 mM NaCl) for 30 days. Scale bars = 1 cm.

Phenotypic segregations in experiments carried out both in vitro and in vivo were consistent with a monogenic recessive inheritance of the mutations (**Supplementary Tables S3 and S4**). Subsequently, a co-segregation analysis was carried out *in vitro* by studying the association between the mutant phenotypes and the *nptII* gene expression. In both mutants, the detection of mutant plants sensitive to kanamycin showed the absence of co-segregation with a functional *nptII* gene (**Supplementary Table S5**).

Finally, a complementation test was performed to determine if the two recessive mutations were alleles of the same gene or of different genes. The test indicated that the two recessive mutations were allelic as they failed to complement each other in the *sga1* x *sga2* hybrid (**Fig. 2A**).

**Fig. 2.**
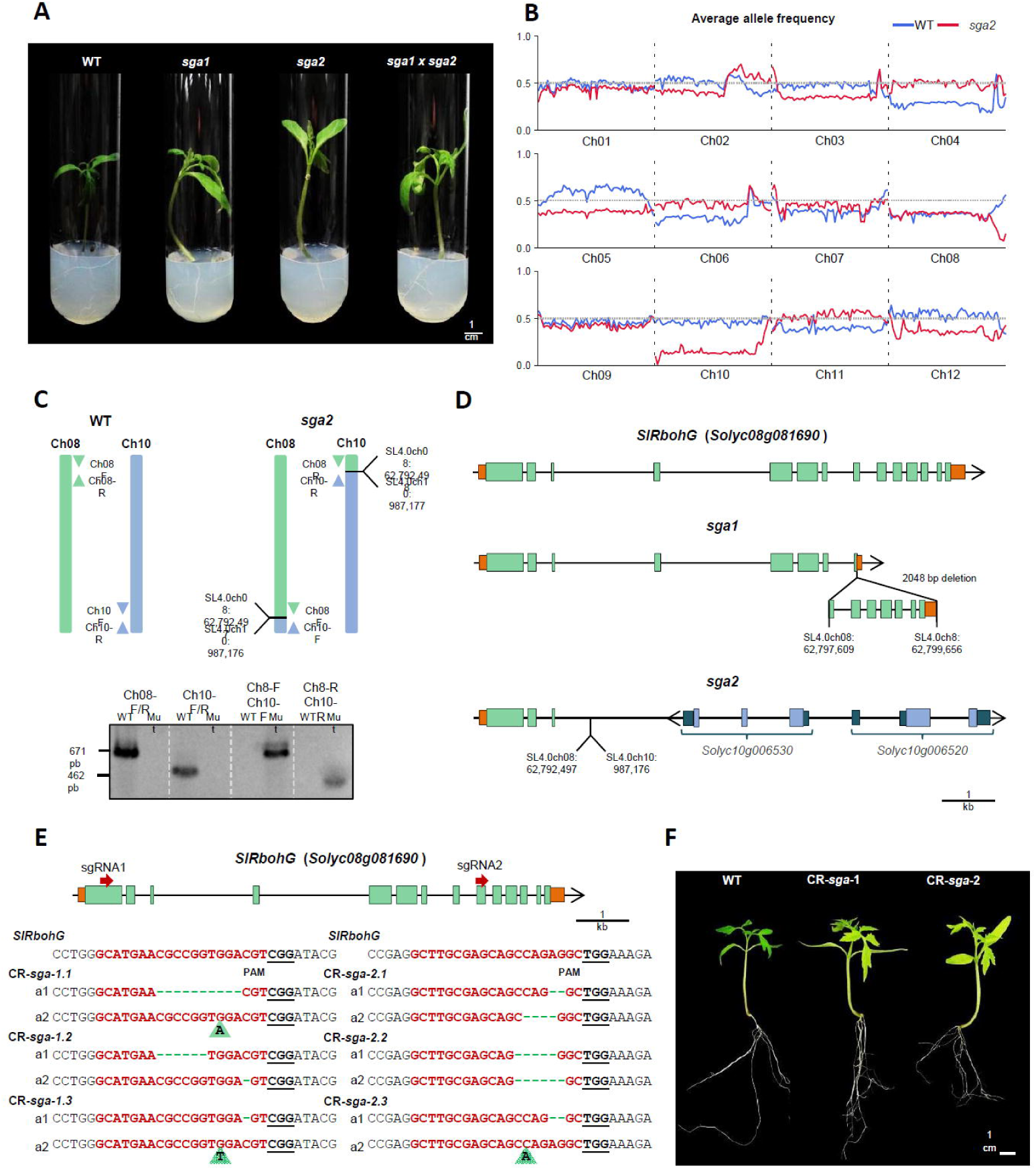
*sga* mutations are allelic and affect the *SlRbohG* gene. A, Allelism test of *sga1* and *sga2* mutants. The F_1_-cross shows both the chlorosis and hyperhydration phenotype of the parent plants (*sga1* and *sga2* mutants). B, Distribution of the average allele frequency of wild-type (WT, blue line) and *sga2* (red line) pools grouped by chromosomes. C, Schematic representation (up) and PCR (down) showing the balanced translocation between chromosomes 8 and 10 found in the *sga2* mutant. D, Schematic representation of the *SlRbohG* gene and the genomic rearrangements resulting from the *sga1* and *sga2* mutations. *SlRbohG* (*Solyc08g081690*) gene coding and UTR sequences are represented by green and dark green boxes, respectively. Coding and UTR sequences of chromosome 10 genes (*Solyc10g006530* and *Solyc10g006520*) are depicted by blue and dark blue boxes, respectively. E, CRISPR/Cas9 alleles identified by cloning and sequencing PCR products from knockout lines using two independent sgRNAs targeting the *SlRbohG* exon 1 (sgRNA1) or exon 9 (sgRNA2). A total of six independent CRISPR/Cas9 diploid lines were molecularly assessed, three for each sgRNA: sgRNA1 (CR-*sga*-1) and sgRNA2 (CR-*sga*-2). Green dashed lines show InDel mutations, green triangles indicate an insertion nucleotide, and black bold and underlined letters indicate protospacer adjacent motif (PAM) sequences. F, Representative sensitive phenotype of CR-*sga* −1 and CR-*sga* −2 lines under salt stress conditions.

### The *sga* phenotypes are due to mutations in the coding sequence of the *SlRbohG* gene

As a T-DNA insertion was not associated with the mutant phenotype observed in *sga1* and *sga2*, a mapping-by-sequencing strategy (**Yuste-Lisbona *et al*. 2021**) was performed to identify the causative mutations using an F_2_ population derived from crossing *sga2* to the wild tomato *S. pimpinellifolium* (LA1589 accession). A total of 103 F_2_ plants were evaluated in an *in vitro* assay under salt stress conditions (100 mM NaCl), from which 25 showed the *sga* phenotype. The phenotypic segregation observed in this interspecific F_2_ progeny (78 WT: 25 *sga*) was consistent with a monogenic recessive inheritance pattern of the *sga2* mutation (χ2 = 0.03, 90% ≥ P ≥ 75%).

Genome sequencing was conducted on two DNA pools containing 30 WT and 25 *sga* F_2_ plants. Genome-wide analysis of the average allele frequencies in the WT and mutant pools revealed two genomic regions on chromosome 8 and 10 with a strong bias towards tomato reference alleles (**Fig. 2B**), indicating that they are candidates to harbor the genetic variant underlying the *sga2* mutation. The examination of genomic paired-end reads in these candidate regions showed the presence of a balanced translocation in the mutant pool, which involves the ending and starting regions of chromosome 8 and 10, respectively (**Fig. 2C**). The translocation breakpoint on chromosome 8 (SL4.0ch08:62,792,497) disrupted the third intron of the *SlRbohG* (*Solyc08g081690*) gene (**Fig. 2D**), whereas the breakpoint on chromosome 10 (SL4.0ch10:987,176) was located in the promoter region (1063 bp from transcriptional start site) of the *Solyc10g006540* gene encoding a formin-like protein. PCR analyses with primers flanking the translocation breakpoints confirmed the presence of this balance translocation in *sga2* (**Fig. 2C**). Co-segregation analysis performed in the F_2_ mapping population showed that all 25 plants with *sga* phenotype were homozygous for the translocation breakpoint on chromosome 8. In contrast, 7 of the 25 F_2_ plants with *sga* phenotype did not bear the chromosome 10 translocation, indicating that the disruption of the *SlRbohG* gene is responsible for the *sga* phenotype. This hypothesis was further validated with the sequencing of the *SlRbohG* locus in the *sga1* mutant, which led to the identification of a 2048 bp deletion (SL4.0ch08:62,797,609..62,799,656) encompassing exons 8-14 and flanking intronic sequence of this gene (**Fig. 2D**).

To demonstrate whether the loss-of-function of *SlRbohG* is responsible for the *sga* phenotype, two different approaches were performed to generate CRISPR/Cas9 knockout lines by targeting the *SlRbohG* exon 1 (sgRNA1) or exon 9 (sgRNA2). A total of six independent CRISPR/Cas9 diploid lines (CR-*Slrbohg*), three for each sgRNA, were molecularly assessed, and all of them were biallelic for edited knockout alleles (**Fig. 2E**). These T_0_ plants were selfed to generate the T_1_ populations, which were evaluated under salt stress conditions. In all cases, CR-*Slrbohg* lines displayed a salt sensitivity phenotype that clearly resembled the one observed in *sga* mutants (**Fig. 2F**). Hence, results revealed that the reported translocation and deletion mutations in the *SlRbohG* gene were responsible for the sensitive phenotype observed in the *sga* mutants under salt stress conditions.

### The disruption of *SlRbohG* gene promotes high shoot water and Na^+^ accumulation under *in vitro* and *in vivo* salt stress conditions

A first *in vitro* experiment was carried out with WT and the *sga1* mutant by applying 75 mM NaCl for 15 days, in which the mutant phenotype clearly showed the hyperhydrated aspect under salt stress. In fact, water content increased up to 70% in mutant compared with WT, while shoot dry weight was significantly reduced (**Supplementary Fig. S3A**). These differences in hydration levels resulted in similar shoot fresh weight in WT and mutant. The experiment was repeated in the same conditions by using both *sga* mutants. In this assay, the *SlRbohG* expression was analyzed in roots, which increased 25% in salt stress in WT (**Supplementary Fig. S3B**). As expected, the amounts of transcripts were undetectable in *sga1* and *sga2* mutants. Another feature of the mutant phenotypes in vitro was the chlorosis induced by salt stress, which could be due to an alteration of the Na^+^ homeostasis. Despite the very low transpiration rate under *in vitro* culture conditions, the Na^+^ content increased by more than 50% in both mutants with respect to WT after 15 DST, and the Na^+^/K^+^ ratio was significantly higher in the mutants than in the WT, since no differences in K^+^ accumulation were observed (**Supplementary Fig. S3C-E)**. Thus, the increased chlorosis, and hence salt hypersensitivity, is due to increased Na^+^ accumulation *in vitro*.

Subsequently, both *sga* mutants were characterized under transpiring conditions *in vivo*. For this purpose, WT and *sga1-2* mutant plants were grown in hydroponic culture, and salt treated (100 mM of NaCl) at the 6^th^ leaf stage for 7 DST. Under these conditions where Na^+^ uptake by plants is faster, the detrimental salt effects in mutants were earlier observed, so after 2 DST mutants showed evident chlorosis and light hyperhydration of leaves symptoms which were more pronounced after 7 DST (**Fig. 3A**). The expression pattern of *SlRbohG* induced by salinity was analyzed both at spatial and temporal levels (**Fig. 3B**). After 2 DST, the highest levels of SlRbohG expression was detected in the root, followed by the adult parts of the shoot (leaf and stem), while no increases were observed in the upper young part of the shoot. Transcripts of *SlRbohG* reached the maximum levels after 2DST, while after 7 DST *SlRbohG* expression decreased to the basal levels found in the absence of salt stress (**Fig. 3B**).The salt treatment caused an early arrest of shoot growth of both mutants only after 2 DST, which was maintained along the treatment (**Fig. 3C**). As expected, significant chlorophyll reduction was observed after 2 DST and continued decreasing at the end of the assay (7 DST) (**Fig. 3C**). While in roots there were no differences in water content between WT and mutants, it increased in leaves of both *sga1-2* mutants after 2 and 7 DST (25% and 50%, respectively) (**Fig. 3D**). Regarding traits related to leaf water transpiration (**Fig. 3E**), there were important reductions of g_s_ and E just after applying salt treatment in WT leaf, which were maintained through 7 DST. The tendency was similar in both mutants, although the reductions were lower in *sga1-2* than WT at 2 and 4 DST but not at 7 DST. In order to deep in different transpirational features between mutants and WT, a similar hydroponic assay was performed with WT and *sga1* mutant plants. Before applying salt treatment, measurements of leaf water loss using detached leaflets from the 2^nd^ leaf were taken (**Supplementary Fig. S4A**). The similar values throughout the period after leaf detachment (48 h) indicated that water lost by transpiration was not faster in the mutant than in WT. Similarly to the previous assay, the leaf water content was 50% higher in *sga-1* than in WT after 7 DST (**Supplementary Fig. S4B).** However, a significant reduction in foliar ψ_s_ was observed in *sga1* mutant with respect to WT after 2 DST and, especially, after 7 DST (**Supplementary Fig. S4C).**

**Fig. 3.**
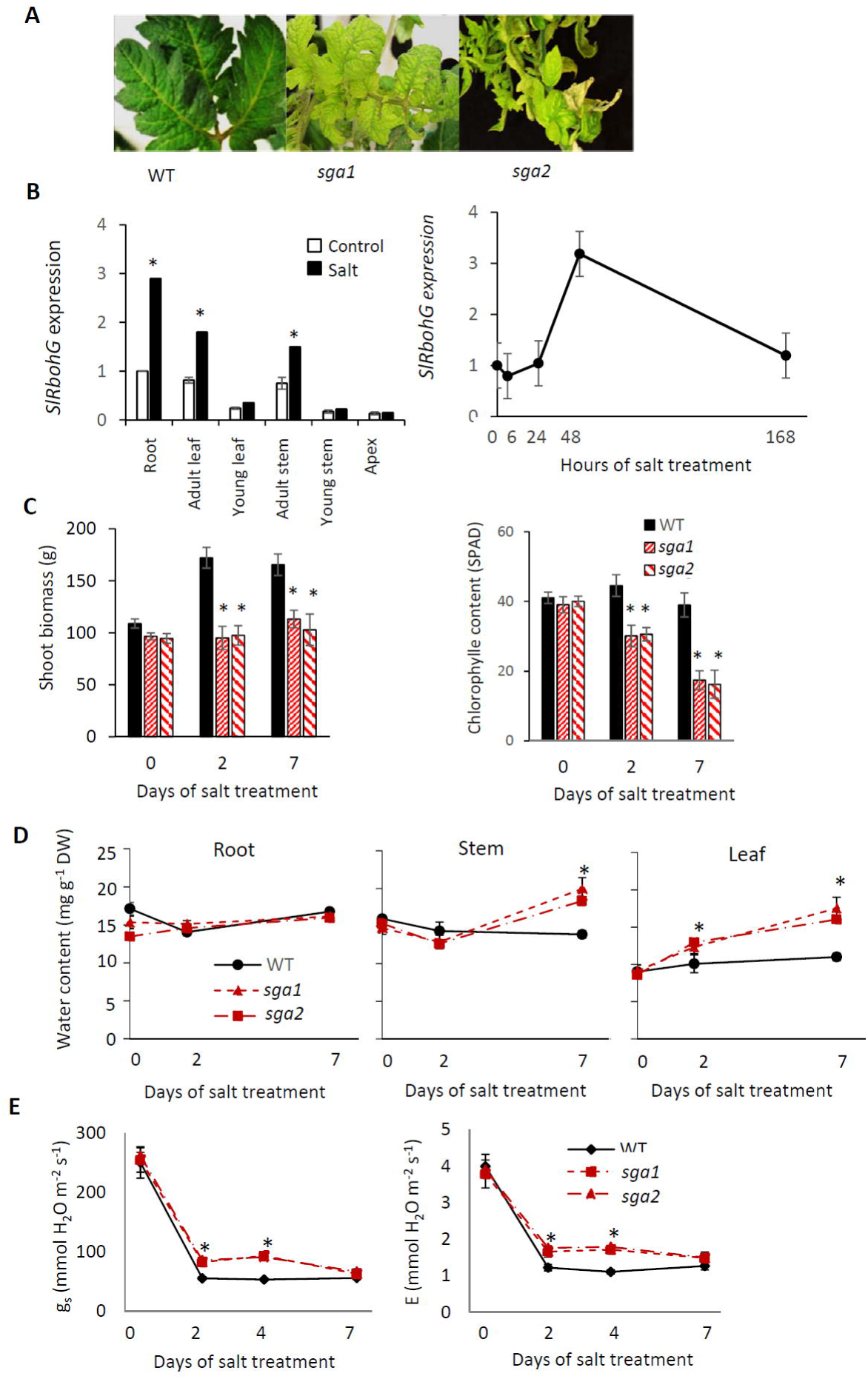
Changes induced by salt stress in *sga1* and *sga2* mutants under *in vivo* transpiring conditions. Plants of WT and *sga1* and *sga2* mutants were grown in hydroponic culture, where the salt sensitivity effects are early observed, and at the 6^th^ leaf stage salt treatment (100 mM NaCl) was applied for 7 days (DST). A, Detail of the salt-treated leaves showing the mutants a high chlorosis degree after 7 DST. B, Spatial expression of *SlRbohG* in root and different shoot parts of the WT plants grown in control and salt for 2 days, and temporal expression of *SlRbohG* in WT root for 7 DST. C, Shoot biomass and chlorophylls in leaves of WT and mutant plants at 0, 2 and 7 DST. D, Water content evolution in adult leaf (2^nd^ leaf), stem (first internode) and root. E, Stomatal conductance (g_s_) and transpiration rate (E) measured in the 2^nd^ leaf throughout salt treatment period. Values are means ± SE of three biological replicates of three plants each. Asterisks indicate statistically significant differences according to Tukey’s test (*p* < 0.05).

As previously showed, one of the main phenotypic symptoms induced by salinity in *sga1-2* mutants was leaf chlorosis, which was related to Na^+^ homeostasis. While the Na^+^contents did not change with respect to WT in roots, the Na^+^ accumulation increased in stems and, especially, in leaves of both mutants during salt treatment period, being 3 times higher in leaves of *sga1-2* mutants than WT after 7 DST (**Fig. 4).** Regarding K^+^, the most significant changes were observed in the root at 2 DST, where K^+^ content was reduced by more than 50% in both mutants with respect to WT (**Fig. 4)**. Interestingly, a key trait related to salt sensitivity, Na^+^/K^+^ ratio, presented increased values in the three organs of both mutants, especially in leaves (**Fig. 4)**.

**Fig. 4.**
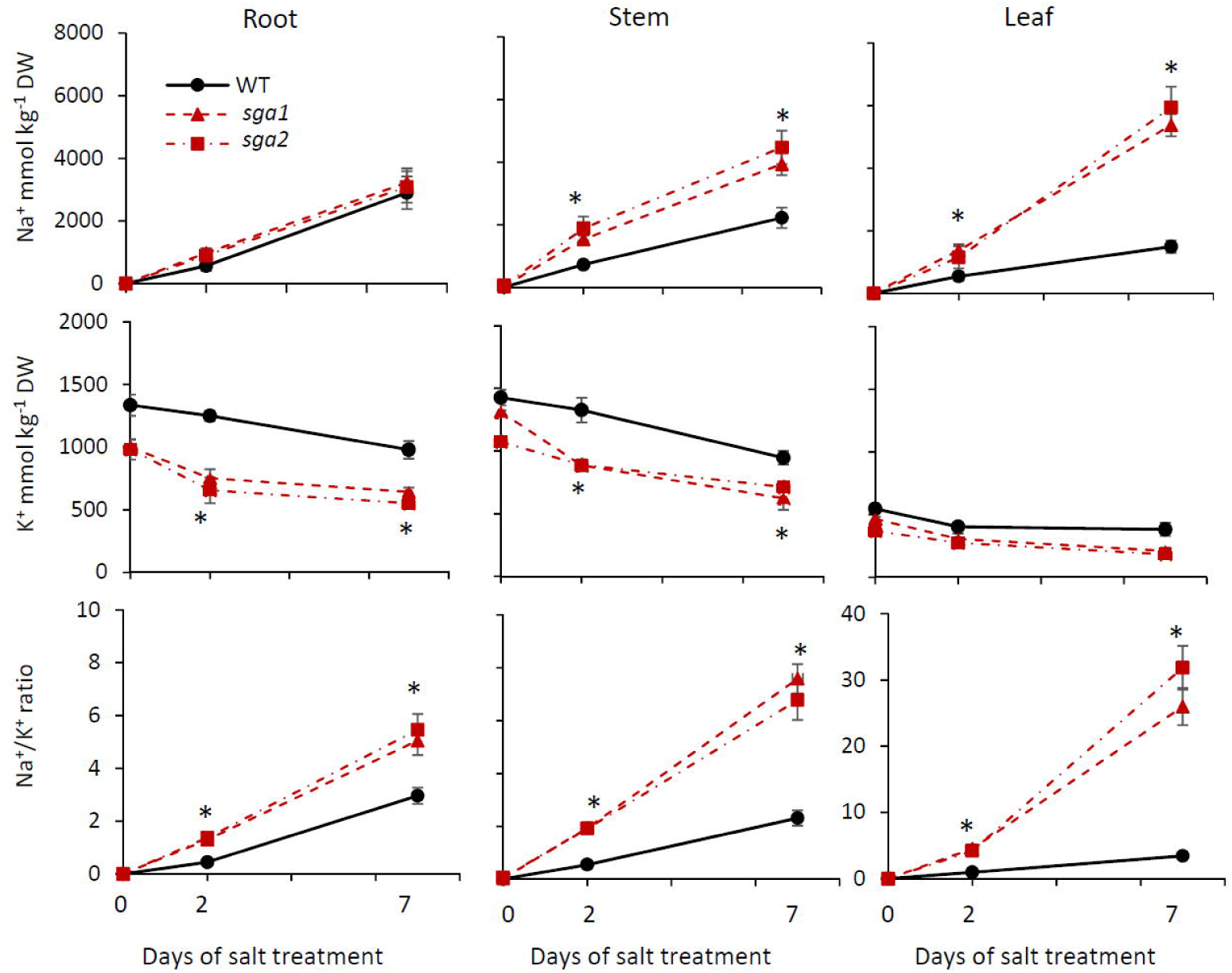
The disruption of *SlRbohG* promotes a massive transport of Na^+^ from root to shoot under *in vivo* transpiring conditions of salt stress. Plants of WT and *sga1* and *sga2* mutants were grown in hydroponic culture, and at the 6^th^ leaf stage salt treatment (100 mM NaCl) was applied for 7 days (DST). Evolution of Na^+^ and K^+^ contents were analyzed in root, stem (intersection of 1st-2^nd^ leaf), and leaf (2^nd^), and Na^+^ /K^+^ ratios were calculated. Values are means ± SE of three biological replicates of three plants each. Asterisks indicate statistically significant differences according to Tukey’s test (*p* < 0.05).

In order to deep in changes induced by salinity in Na^+^ distribution along mutant plants the experiment was repeated by using WT and *sga1* mutant, and the Na^+^ pattern was analyzed in the root, basal and upper stems and leaves as well as young (developing) leaves (**Supplementary Fig. S5**). Results showed that the mutant accumulated higher Na^+^ content than WT in all vegetative tissues analysed, both in young apical tissues as well as more basal and older tissues, except in root, where the Na^+^ content was similar throughout the treatment.

### The expression pattern of genes involved in Na^+^ homeostasis and water transport is altered in *sga* mutants under salinity

Along with chlorosis, another symptom induced by salt stress in both mutants was their hyperhydration. Since AQPs are involved in water transport, we analysed the expression levels of 6 isoforms of plasma membrane aquaporins (*SlPIP1;3*, *SlPIP1;5*, *SlPIP1;7*, *SlPIP2;1*, *SlPIP2;10*, *SlPIP2;12*) and 2 isoforms of tonoplast (*SlTIP1;1* and *SlTIP2;2*) in roots and leaves of WT and *sga1-2* mutants grown in control and after 2 DST. Firstly, we identified the basal expression levels of isoforms in WT and mutant plants, observing that the isoform exhibiting the highest expression in both root and leaf is *SlPIP1;3*, followed by *SlPIP1;5* and *SlPIP2;1* (**Fig. 5A**). The changes induced by salinity in the expression levels of AQPs genes in WT were observed mainly in leaves, where almost all AQPs analysed increased their expression, mainly of *SlPIP1;3*, *SlPIP1;5* (**Supplementary Fig. S6**). Contrary, the expression of *SlPIP2;12* was strongly inhibited by salinity in leaves and mainly in root of WT (**Supplementary Fig. S6**). Curiously, in the root of *sga* mutants, *SlPIP2;12* gene was the only AQPs with a significantly increased expression than in WT, while the rest AQPs showed a similar expression level. In leaves, compared to WT plants, none of the AQPs showed an increased level, but also some of them were reduced with respect to WT, as observed for SlPIP1;7, SlPIP2;1 and SlPIP2;10 (**Fig. 5B**).

**Fig. 5.**
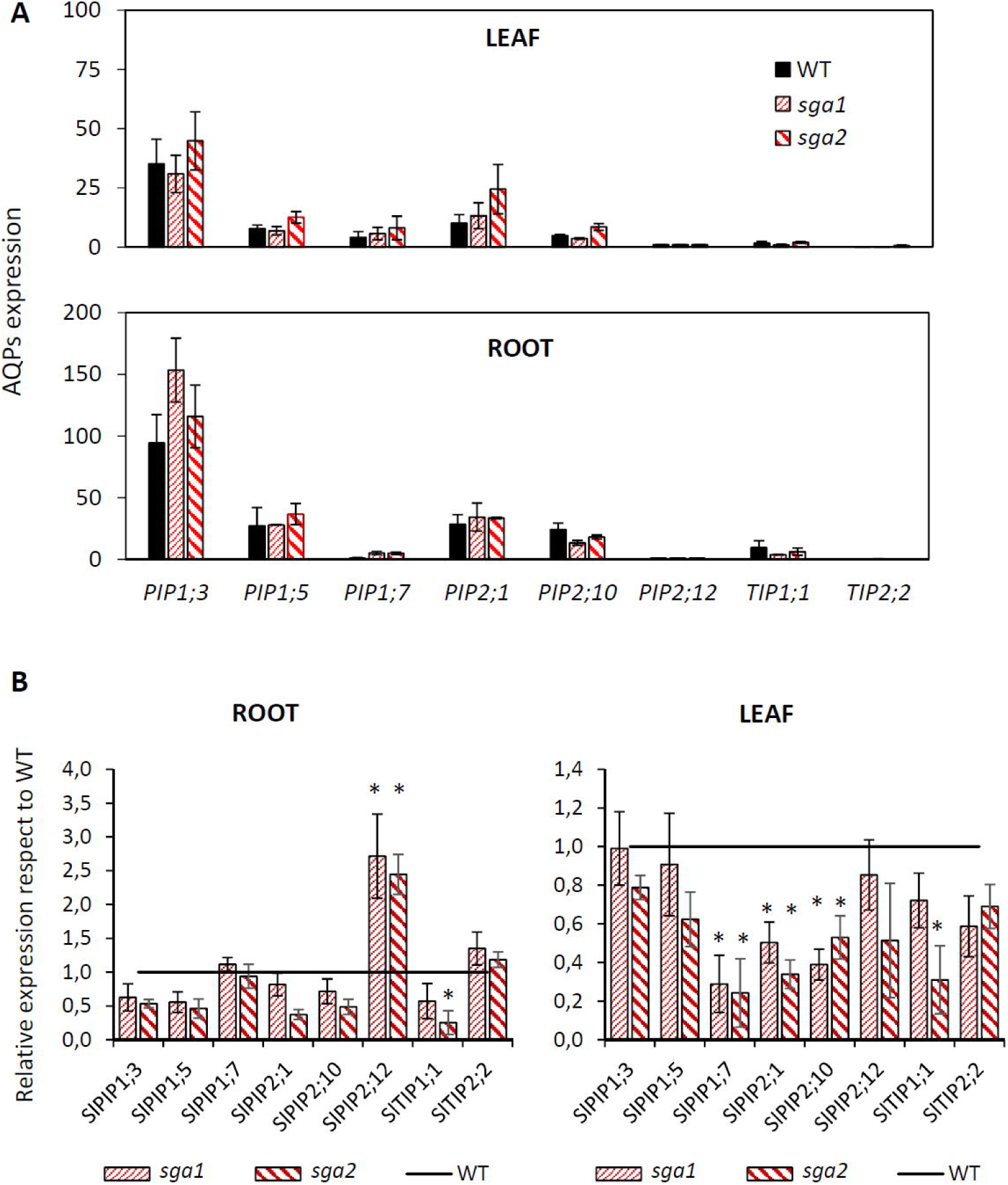
Expression levels of aquaporins genes in WT and *sga1* and *sga2* mutants. Relative expression of six genes of *SlPIP* subfamily and two of *SlTIP* subfamily were analyzed in root and leaf (2^nd^ leaf) prior to salt stress (control) and after 2 days of 100 mM NaCl treatment. A, Relative expression of different isoforms in WT and *sga* mutants under control conditions. The expression of *SlPIP2;12* in WT was set to 1. B, Expression changes induced by salt in both mutants with respect to WT. For each gene, the expression in WT was set to 1. Values are means ± SE of three biological replicates of three plants each. Asterisks indicate statistically significant differences with respect WT according to Tukey’s test (*p* < 0.05).

Since both mutants are characterized by high transport of Na^+^ from root to shoot, the expression levels of genes involved in Na^+^ transport *(SlSOS1, SlHKT1;1* and *SlHKT1;2*) and compartmentation in the vacuole (*SlNHX3* and *SlNHX4*) were analyzed in roots of WT and *sag1-2* mutants grown in control and after 2 DST (**Fig. 6A**). The expression of *SlSOS1* and specially *SlHKT1;1* and *SlHKT1;2* was strongly down-regulated in roots of the *sag* mutants under control and salt stress. With respect to genes involved in vacuolar compartmentation, the expression was only reduced in *SlNHX3* in both control and salt, while *SlNHX4* was only reduced in control. In addition, and as a consequence of K^+^ homeostasis alteration, the genes involved in K^+^ homeostasis *SlHAK5* and *SlSKOR* were also analyzed (**Fig. 6B**). While there were no differences between WT and mutants for *SlHAK5*, the *SlSKOR* expression was significantly reduced in both mutants with respect to WT roots.

**Fig. 6.**
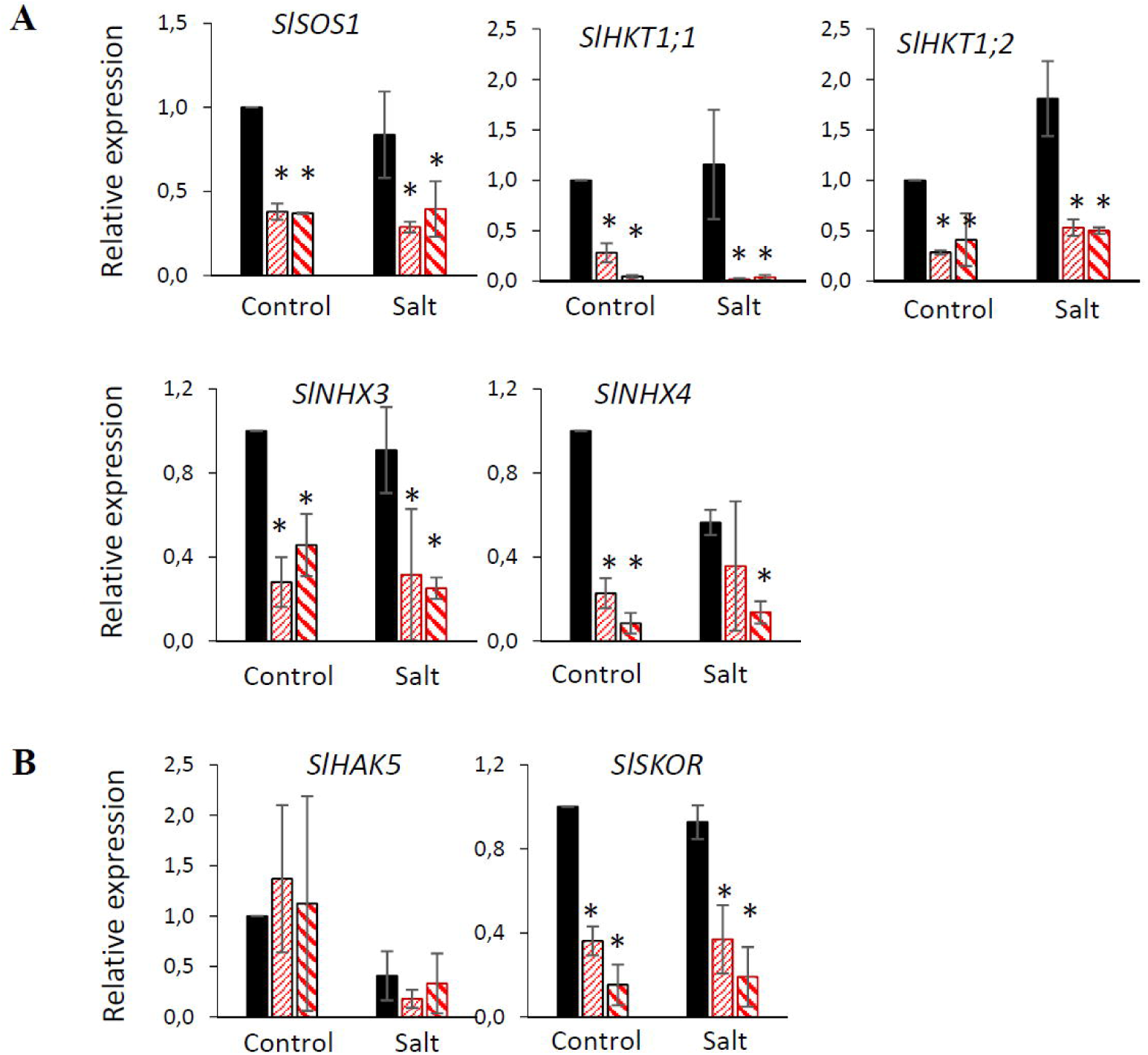
Disruption of *SlRbohG* downregulates the genes expression involved in Na^+^/K^+^ homeostasis. Genes were analyzed in root prior to salt stress (Control) and after 2 days of 100 mM NaCl treatment (Salt). A, Relative expression of genes involved in Na^+^ uptake and transport from root to shoot (*SlSOS1, SlHKT1;1* and *SlHKT1;2*), and vacuole compartmentation (*SlNHX3* and *SlNHX4*). B, Relative expression of genes involved in K^+^ uptake and transport. The expression of genes in WT at day 0 was set to 1. Values are means ± SE of three biological replicates of three plants each. Asterisks indicate statistically significant differences respect WT according to Student’s test (*p* < 0.05).

### The *SlRbohG* gene disruption alters the H_2_O_2_ production

*SlRbohG* expression induced by salt stress is generally associated with increased H_2_O_2_. In order to know the changes caused by the mutation in this aspect, H_2_O_2_ production was analyzed by means of *in vitro* and *in vivo* assays. *In vitro*, a higher fluorescence level and increased production of H_2_O_2_ was found in roots of salt-treated WT seedlings for 15 days, while the H_2_O_2_ production did not change with salt stress in *sga* mutants (**Fig. 7A**). Next, we analyzed the H_2_O_2_ production of the plants grown *in vivo* (**Fig. 7B**). A higher fluorescence level was only observed in WT plants after 2 DST, which was corroborated when the H_2_O_2_ content was measured by spectrophotometry (7 DST). Thus, the H_2_O_2_ production increased with salinity in the three plant parts of WT, while no changes were observed in *sga1*-*2* mutants. Taken together, the increased expression of *SlRbohG* was associated with H_2_O_2_ accumulation in WT plants, and this response was similar in both *in vitro* and *in vivo* conditions.

**Fig. 7.**
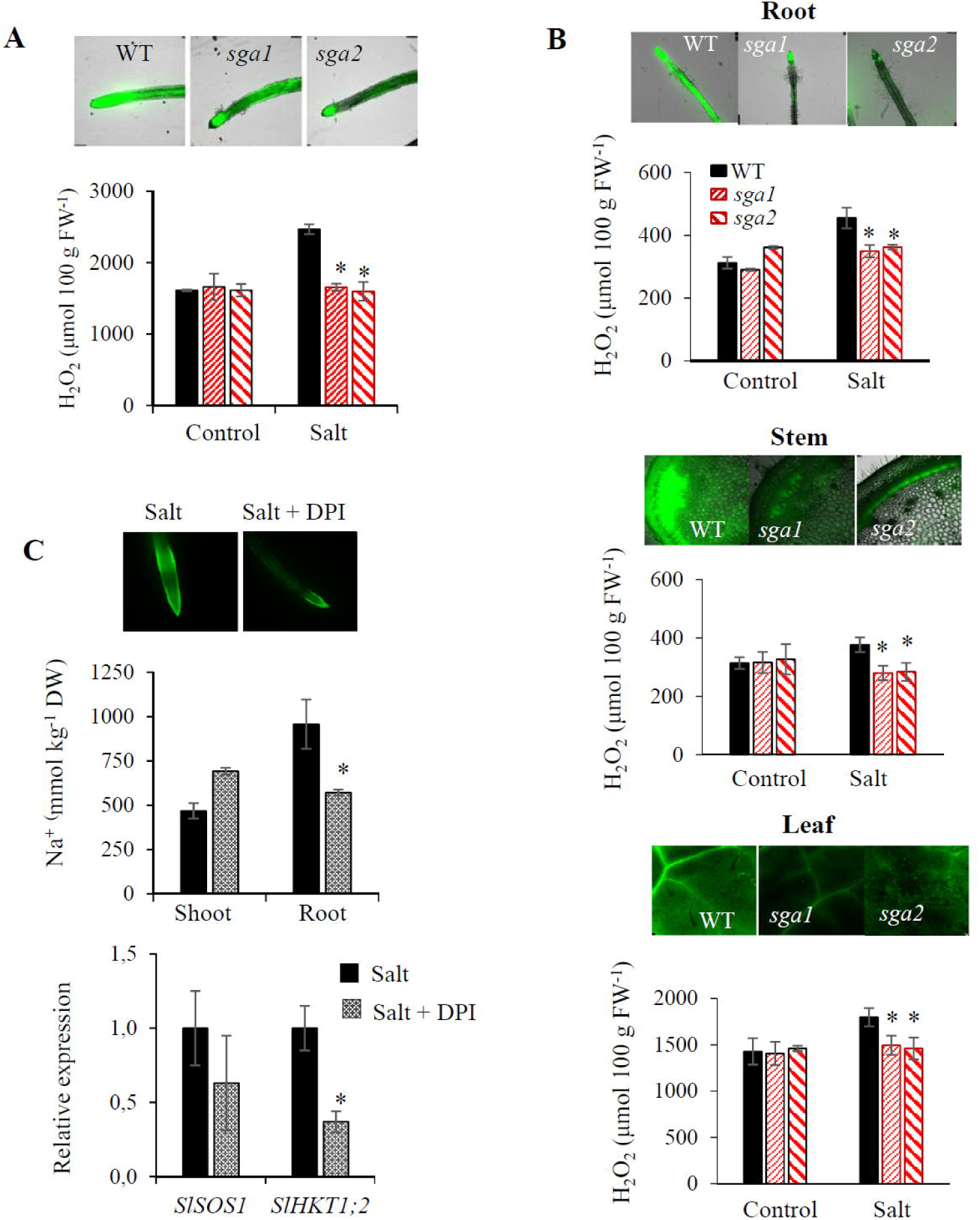
H_2_O_2_ production under *in vitro* and *in vivo* conditions. H_2_O_2_ was analyzed by confocal microscopy and by spectrophotometry using e-FOX assay. A, *In vitro*, WT, *sga1* and *sga2* plants were grown under control and salt stress (75 mM NaCl) for 15 days. H_2_O_2_ concentration was measured in roots at the end of the experiment. B, *In vivo*, WT, *sga1* and *sga2* mutants were grown in a hydroponic system and salt stress (100 mM NaCl) was added to the nutrient solution at the 6^th^ leaf stage for 7 days (DST). H_2_O_2_ concentrations were measured in roots, stem (between 1^st^ and 2^nd^ true leaves) and leaf (2^nd^ leaf) of control and salt-treated plants after 2 DST (microscopy) and 7 DST (spectrophotometry). C, Effects of NADPH oxidase inhibitor diphenylene iodonium (DPI) on H_2_O_2_, Na^+^ accumulation and Na^+^ transporter expression in roots of WT plants grown in control conditions and treated with salt stress (75 mM NaCl) alone or adding DPI inhibitor (100 μM) at the 2-leaf stage. Shoot and root Na^+^ concentrations were measured after 2 days of salt and DPI treatments, and H_2_O_2_ accumulation and *SlSOS1* and *SlHKT1;2* relative expression were measured in roots after 1 day. Values are the mean ± SE of two independent assays. For *in vitro* experiments, at least six independent samples per treatment were assayed with two seedlings (two developed leaves stage) per sample. For *in vivo* assays, plants of 6-7 fully expanded leaves separated into three different tissues (first-developed leaflets, central stem sections and longitudinal roots samples) were used. Asterisks indicate statistically significant differences respect WT according to Tukey’s test (*p* < 0.05).

### The treatment of WT with the NADPH-oxidase inhibitor DPI increases the Na^+^content in the shoot due to reduced expression of *SlHKT1;2* in the root

To corroborate previous results, we studied in salt-treated WT plants the effects of NADPH oxidase inhibitor diphenylene iodonium (DPI) on H_2_O_2_, the Na^+^ levels and the amounts of *SlSOS1* and *SlHKT1;2* transcripts in root (**Fig. 7C**). The H_2_O_2_ fluorescence was lower in WT salt-treated roots with DPI than without DPI inhibitor. The Na^+^ contents increased with DPI treatment in shoots while decreased in roots. Finally, the *SlSOS1* expression did not change with DPI treatment in root, while the *SlHKT1;2* expression level was reduced by 60%. These results suggest that the inhibition of NADPH oxidases in WT salt-treated plants has a similar effect to the disruption of *RBOHG* gene in *sga1-2* mutants, as the H_2_O_2_ reduction in root is associated with higher Na^+^ transport to the shoot via reduced expression of *SlHKT1;2*.

### *SlRbohG* primarily executes its role in response to salt stress from the roots

Until now, we have shown that the salt hypersensitive phenotype of *sga1-2* mutants is caused by a massive transport of Na^+^ to the shoot as mutants can not retain Na^+^ in root. To advance in the study of *SlRbohG* role in root and its contribution to salt stress tolerance, a first grafting experiment was carried out in pots with adult plants by using reciprocal grafting between WT and *sga1-2* mutants and self-grafting plants as reference. After 15 days of 100 mM of NaCl, the salt sensitive phenotype only appeared when the mutants were used as rootstock but not as scion (**Supplementary Fig. S7**).

Next, WT and *sga1* mutant plants were grown without salt (control) and with 100 mM NaCl for 5 days under hydroponic culture because salt effects are observed early.The phenotypes of all grafted combinations were similar under control conditions. Under saline conditions, the hypersensitivity phenotype was only observed when the *sga1* mutant rootstock was used. In contrast, no hypersensitivity phenotype was observed with a WT rootstock and a scion of the *sga1* mutant. (**Fig. 8A, B**). A higher Na^+^ accumulation and Na^+^/K^+^ ratio were observed in phenotypes exhibiting the salt hypersensitivity phenotype (**Fig. 8C**), suggesting that the *SlRbohG* gene primarily plays its role in roots.

**Fig. 8.**
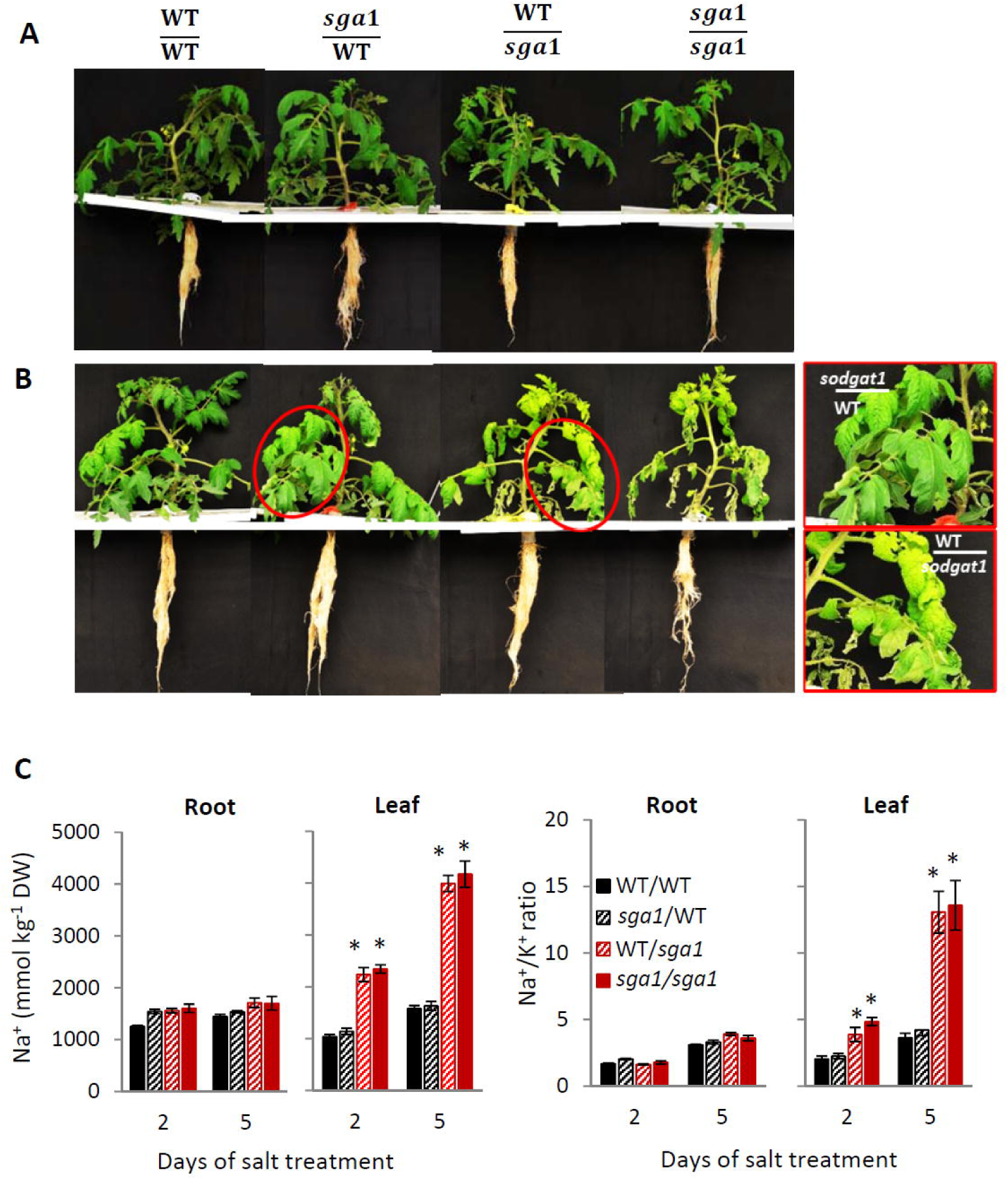
The root is the main plant organ responsible of the salt sensitivity caused by disruption of *SlRbohG* gene. Self-grafting plants of WT and *sga1* mutant (WT/WT and *sga1/sga1*) and reciprocal grafting between WT and mutant were grown in hydroponic culture and at the 6^th^ leaf stage 100 mM NaCl treatment was applied for 5 days. A, The phenotypes are similar in all graft combinations before applying the salt treatment. B, After 5 days of salt treatment the graft combinations where the mutant is used as rootstock show salt sensitivity. At right, detail of the salt-treated leaves of reciprocal grafting. C, Concentrations of Na^+^ and Na^+^ /K^+^ ratio in roots and leaves (2^nd^ leaf) after 2 and 5 days of salt treatment. Values are means ± SE of three biological replicates of three plant each. Asterisks indicate statistically significant differences between mean values of WT/WT and each graft combination according to Tukey’s test (*p* < 0.05).

### The salt sensitivity caused by the disruption of *SlRbohG* also affects reproductive organs

In order to know whether the loss of *SlRbohG* function induces salt sensitivity to long-term, a first assay was performed with adult plants of WT and *sodgat1* plants grown in the same pot, and a high salt level (200 mM NaCl) was applied when the plants had 10^th^-true leaves. The salt effect on the chlorosis provoked in the leaves was quick, as at only 4 DST, the leaf chlorosis of *sga1* was very evident (**Supplementary Fig. S8**). This negative effect continued increasing with the treatment period, with senescent leaves after 12 DST and dead plants after 26 DST.

Finally, a long-term assay was performed in the greenhouse, where the plants were grown at optimal growing conditions (Control) or under salt stress conditions (100 mM NaCl. The salt sensitivity phenotype was corroborated, as the plants died after 40 days of 100 mM NaCl (**Fig. 9A**). At the reproductive level, the sga1 plants did not produce fruit, which was due to the fact that the flowers started to dry early and the trusses could not develop like those of the WT plants. In order to link this phenomenon with the massive influx of toxic Na^+^ into reproductive organs, the content of this ion was analyzed in anthesis flower and green fruit of WT and *sga1* mutant (**Fig. 9B**). The results confirmed a notably higher concentration of Na^+^ in both the flower and fruit of the mutants compared to WT. Altogheter, results indicate that the disruption of *SlRbohG* provokes high salt sensitivity at long-term and at both levels, vegetative and reproductive.

**Fig. 9.**
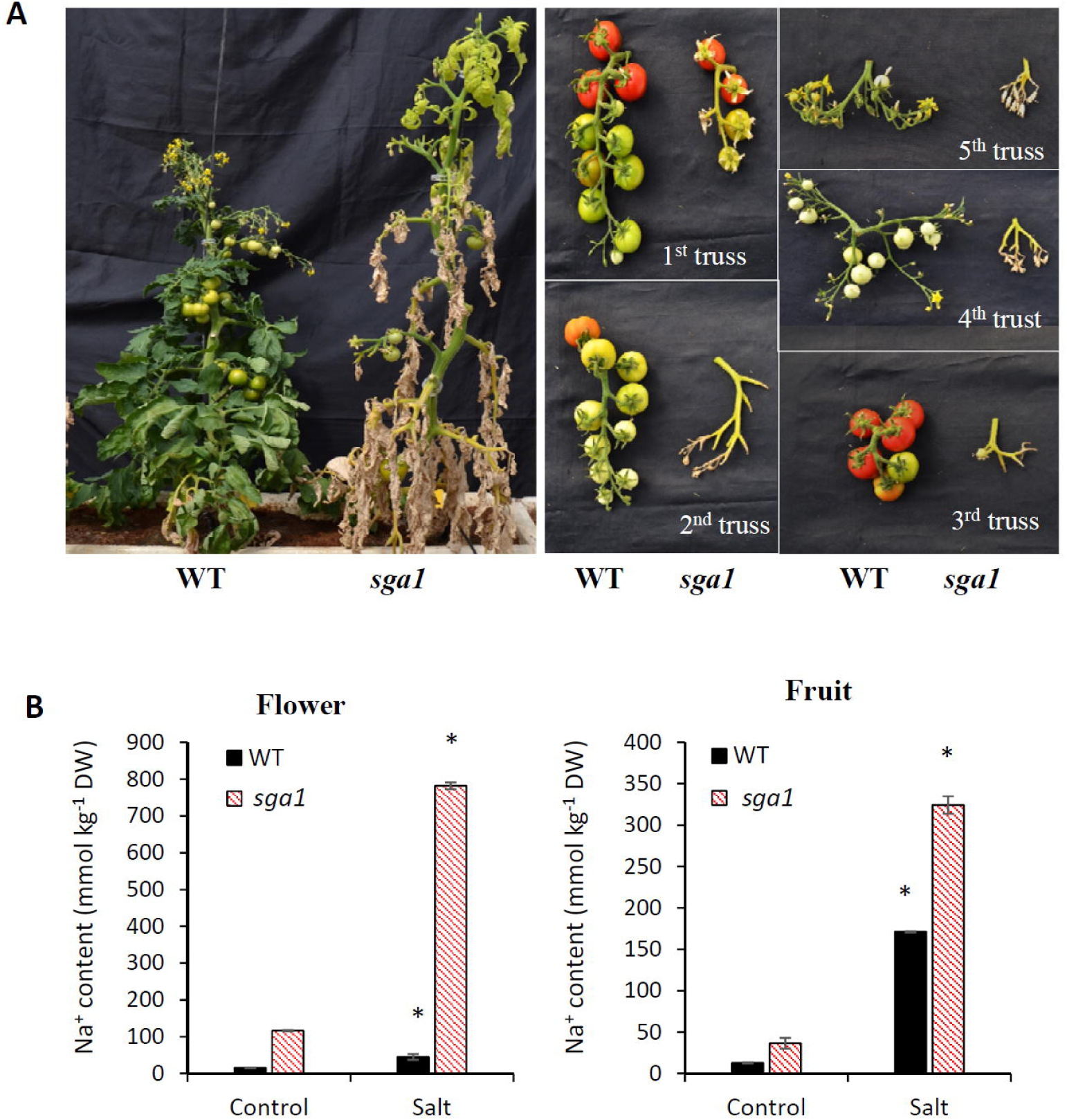
The *SlRbohG* function is essential for tomato reproductive development under salt stress. WT and *sga1* plants were grown in a greenhouse during spring-summer season, and when the plants had 8 true leaves salt treatment (100 mM NaCl) was applied. A, Photographs representing entire plants and trusses from the first to the fifth after a 40-day salt treatment B, Na^+^ content was analyzed in anthesis flowers and green fruits of WT and sga1 mutant after 40 days under control and salt treatment conditions. Values are means ± SE of three biological replicates of three plants each. Asterisks indicate statistically significant differences with respect WT at the same condition according to Tukey’s test (p < 0.05).

To further investigate the cause of aerial tissues of *sga* mutants swelling due to water accumulation under salinity conditions, microscopic analysis of the degree of lignification of the *sga1* mutant was performed. These studies confirmed that the degree of lignification of vascular bundles in the petioles of *sga1* mutant was significantly lower than that of the WT (**Supplementary Fig. S9**). This reduced lignification could provide greater flexibility and permeability to cell walls, positively impacting the transport and retention of water in the cells, which would explain the higher hyperhydration under saline conditions.

## DISCUSSION

Here we present the identification and characterization of two tomato allelic mutants, named *sga1* and *sga2*, that exhibit extreme salt hypersensitivity due to elevated Na^+^ accumulation in shoots. We established that the salt hypersensitivity observed in these mutants results from the loss of function of the *SlRbohG* gene, an ortholog of the Arabidopsis *AtRbohF* gene. While the role of this gene, also known as *Rboh1*, has been involved in plant development and stress response **(Shi *et al*., 2015; Chen *et al*., 2016; Yin *et al*., 2018)**, its fundamental function in regulating salt stress tolerance in tomato has not been elucidated to date. Additionally, the *sga* mutants described herein represent the first identified lines with the *SlRbohG* gene mutated, marking a significant advancement, as prior investigations into its function relied on transgenic lines.

### The expression of the *SlRbohG* gene in roots is crucial to ensure the level of salt tolerance of tomato plants throughout their life cycle

Disruption of *SlRbohG* gene in *sga* mutants promoted a salt-sensitive phenotype characterized by pronounced leaf chlorosis and swelling from the early stages of salt stress, which was observed under both in vitro and in vivo culture conditions. This effect persisted in young as well as adult plants. At the reproductive stage, *sga* mutants experienced a significant reduction in fruit production due to flower abscission or premature dying before fruit setting. Notably, these findings represent the first results revealing the pivotal role of *SlRbohG* gene in the long-term salt tolerance of tomatoes.

To delve into the functions of *SlRbohG* in salt tolerance, we conducted investigations on its role in roots and shoots separately through reciprocal grafting assays between WT and *sga* mutants. The salt-sensitive phenotype was evident only when the mutants served as rootstocks, not as scions, underscoring the root as the primary plant organ responsible for the salt sensitivity induced by the disruption of the *SlRbohG* gene. Thus, the loss of function of the *SlRbohG* gene in tomato roots significantly impairs salt tolerance throughout its entire life cycle, affecting vegetative and reproductive organs.

### The disruption of the *SlRbohG* gene resulted in shoot hyperhydration under salt stress

One physiological feature observed in both *sga* mutants was their higher water accumulation induced by salinity in the shoot, which was observed under both *in vitro* and *in vivo* conditions. Hyperhydration resulting from the *SlRbohG* mutation had not been described previously, neither in transgenic tomato lines nor in the Arabidopsis mutant (*AtRbohF*). Given that the heightened water accumulation in mutants could not be ascribed to altered transpiration, and recognizing the pivotal role of aquaporins (AQPs) in plant water transport, we opted to explore whether such hyperhydration could be associated with a modified expression pattern of six genes coding for PIPs and two coding for TIPs in roots and leaf. The AQPs isoforms was selected within the large multifunctional gene family of tomato based on its previous described induction in response to abiotic stresses (**Sade *et al*., 2009; Jia *et al*., 2020**). These previous studies have revealed notable variations in the expressions of AQP isoforms in both the roots and leaves of tomatoes, indicating a potential role for SlPIPs in mediating water transport under salt stress conditions. Our findings in the current study, under experimental salt stress conditions, showed an upregulation of *SlPIP1;3* and *SlPIP1;5* expression in leaves, coupled with a significant inhibition of *SlPIP2;12* expression in roots of WT plants. Similar downregulation of transcript levels for specific aquaporins in roots in response to salinity has been documented in tomato and Arabidopsis (**Boursiac *et al*., 2005; Jia *et al*., 2020**). By contrast, in *sga* mutants, the *SlPIP2;12* gene expression exhibited a significant increase compared to WT (**Fig. 5**), which could eventually be associated with increased water transport from the root to the shoot.

In addition to AQPs, another potential contributing factor to *sga* mutant hyperhydration is that the *SlRbohG* gene may preserve the function described for its Arabidopsis ortholog (AtRbohF), particularly in relation to the lignification process (**Lee *et al*., 2013**). Lignin deposition, a major modification of the plants’ cell wall, is triggered in response to various environmental stresses, including salt stress. This strengthening of cell walls aids in resisting salt-induced water stress and helps prevent the excessive entry of Na^+^ ions into cells. **Lee *et al*. (2013)** proposed a model in which a subcellular precision in lignin polymerization results from the combined action of locally restricted production of ROS substrate by the NADPH-oxidase encoded by *AtRbohF* gene and localized peroxidase activity. In the current study, we observed that the degree of lignification in the vascular bundles of the petioles in *sga1* mutant was significantly lower than that of the WT, which in turn can also confer greater flexibility and permeability to cell walls, positively affecting the transport of water and nutrients across the cell membrane and cell walls.

### The disruption of *SlRbohG* gene induces high Na^+^ transport from root to shoot

The *sga1-2* mutants were identified *in vitro* and characterized by a high degree of chlorosis in the shoot and increased Na^+^ accumulation. It is known that Na^+^ accumulation in shoots of plants grown under salt stress is dependent on both the transpiration rate and the Na^+^ concentration in the transpiration stream. The *sga1-2* mutant phenotypes should not have been detected *in vitro* (non-transpiring conditions) if the higher root-to-shoot Na^+^ transport caused by disruption of *SlRbohG* gene were due to higher transpiration, as it occurred with other genes involved in transpiration, such as Arabidopsis *AtHKT1;1* (**Davenport *et al*., 2007**) and tomato R1-MYB gene (**Campos *et al*., 2016**). Therefore, the fact of identifying both *sga* mutants under *in vitro* conditions hints at the fact that the most important function of *SlRbohG* is related to the ionic stress as reflected by the Na^+^ toxicity. The Na^+^ transport from root to shoot was higher when we studied the *in vivo* response of both mutants (transpiring conditions), as Na^+^ accumulation increased in the stem and, especially, in leaves after 7 DST. These results indicate that salt sensitivity induced by the *SlRbohG* disruption is mainly due to an alteration of the Na^+^ homeostasis.

The so high Na^+^ transport from root to shoot in the mutants is associated with an altered gene expression pattern of Na^+^ transporters in root. Since *SlSOS1* is involved in root Na^+^ upload to xylem (**Olias *et al*., 2009**), down-regulation of the *SlSOS1* expression reflects that uploading of Na^+^ in root xylem was lower in the *sga* mutant roots. However the high Na^+^ transport from root to shoot was directly related to the intensely down-regulated expression of both *SlHKT1;1* and *SlHKT1;2* in roots of the *sga* mutants. The *SlHKT1;1* and *SlHKT1;2* genes are the main responsible for the removal of sodium from the xylem into the cells in tomato and their wild relatives (**Asins *et al*. 2013**, **Albaladelo *et al*., 2017**; **Egea *et al*., 2018**). Another relevant question involved in Na^+^ homeostasis is the capacity to sequester Na^+^ in vacuoles via increasing expression of *LeNHX3* and *LeNHX4* (**Galvez *et al*., 2012**). *LeNHX3* expression was reduced in roots of both sga mutants, suggesting that in addition to the higher Na^+^ transport up to the leaves, the compartmentation capacity in vacuoles is also lower in *sga1-2* mutants. Regarding K^+^ homeostasis, we observed that the *SlSKOR* channel gene expression was also reduced in the roots of both mutants compared with WT, which suggests that the lower K^+^ transport to the mutant shoot is due to the altered expression of this gene involved in xylem K^+^ uploading (**Nieves-Cordones *et al*., 2014**). In sum, the high shoot Na^+^ accumulation in mutants is related to the high reduction of *HKT1s* expression levels, which impairs the Na^+^ retrieval from the xylem in roots and, as a result, increases Na^+^ transport from root to shoot (**Fig. 10**). Moreover, the altered K^+^ homeostasis in the shoot of *sga* mutants is also related to the altered gene expression pattern of *SKOR* channel in root.

**Fig. 10.**
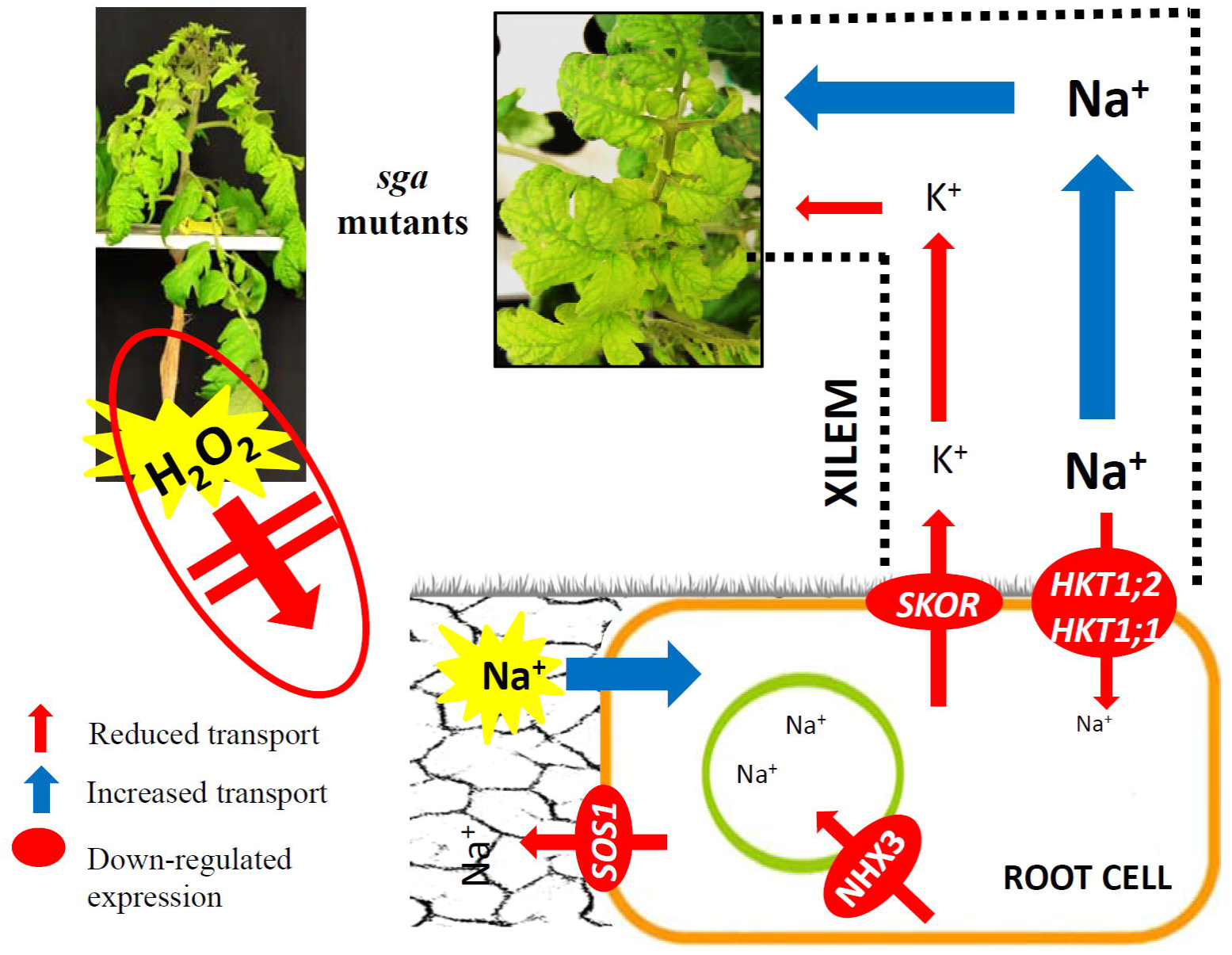
A possibly model of alteration of Na^+^/K^+^ homeostasis in *sga* mutants caused by the lack of H_2_O_2_ production to act as signal molecule and regulate ion homeostasis under salt stress. In root, the intense down-regulation of *SlSOS1* together with *SlHKT1s* induces high uptake and transport of Na^+^ via xylem up to achieve the leaves, which causes leaf Na^+^ toxicity reflected in a high degree of leaf chlorosis. Contrarily, the K^+^ channel is repressed by reducing the K^+^ transport. The thickness of arrow represents the flow size.

One relevant and unsolved question is whether the high water transport from root to shoot is associated with a cotransport water – Na^+^. In this regard, concepts of plant membrane transport have been generally based on the assumption that water and solutes move across membranes via separate pathways. However, in addition to their major role as specific carriers for ions, membrane transporters can also behave as water channels, although there is a great knowledge gap on this issue. **Wegner (2017)** reviewed experimental evidence for a cotransport of water and solutes, in particular via ion channels that provide pathways for both ion and water transport, as exemplified for maxi K^+^ channels from cytoplasmic droplets of *Chara coralline*. In addition to ion channels, other candidates for solutes-water flux coupling have been proposed, as cation-chloride cotransporters of the CCC type, and transporters of sugars and amino acids (**Wegner, 2017**). However, future studies will be necessary to fully elucidate the cotransport Na^+^ - water.

Ultimately, the substantial accumulation of Na^+^ in shoot results in a significantly lower leaf water potential (ψs) in the *sodgat1* mutant compared to the WT. The altered ψs, influenced by the elevated Na^+^ levels, may play a crucial role in shaping the osmotic conditions that drive water movement within the *sga* mutant (**Boursiac *et al*., 2005**).

### Hydrogen peroxide production is critical for salt stress tolerance

H_2_O_2_ acts as a signaling molecule, participating in a series of physiological and biochemical processes (**Baxter *et al*., 2014**), being RBOHs involved in H_2_O_2_ generation (**Yi *et al*., 2015**). In our study, the higher *SlRbohG* expression induced by salt stress in WT was associated with increased H_2_O_2_ production, which did not occur in *sga* mutants. Interestingly, when the effects of RBOH inhibitor DPI were studied in salt-treated WT plants, we observed that the H_2_O_2_ reduction was associated with Na^+^ accumulation that increased in shoot and decreased in roots, due to the reduced expression of *SlHKT1;2*. These results show that inhibition of NADPH oxidases activity in WT salt-treated plants has a similar effect to the disruption of *RBOHG* gene in *sga1-2* mutants. This observation agrees with suggestions from different authors on the importance of H_2_O_2_ as a key signaling molecule involved in regulation of Na^+^ transport under salt stress (**Niu *et al*., 2018; Li *et al*., 2019**). Moreover, an increase in RBOH-dependent H_2_O_2_ production has been proposed as a mechanism to increase HKT-mediated Na^+^ unloading from the xylem sap (**Jiang *et al*., 2013**).

H_2_O_2_ also serves as a signal in controlling water uptake under abiotic stress since it has similar molecular properties to H_2_O and moves across the plasma membrane via AQPs, although the functions of H_2_O_2_ in water uptake are still rather obscure (**Castro *et al*., 2021**). In this study, an inverse relation seems to exist between H_2_O_2_ production and water accumulation under salt stress, as the water content increased in mutants where no increase in H_2_O_2_ production was detected. The question to solve in future studies will be elucidating how H_2_O_2_ is involved in the physiological processes altered by disruption of *SlRbohG*. Our current knowledge on how plant tissues sense salt stress is limited enough to date (**Kudla *et al*., 2018**), and it would be highly interesting to reveal if this gene is acting as a Na^+^ sensor.

In conclusion, the discovery of allelic mutants *sga1* and *sga2*, highlights the function of the *SlRbohG* gene in the root as essential for ensuring salt tolerance throughout the entire life cycle of tomato. We propose a potential model where the increased production of H_2_O_2_ mediated by *SlRbohG* under salt stress conditions is needed for inducing the expression of genes coding for Na^+^ transporters in the roots, particularly *SlHKT1;2*, responsible for withdrawing Na^+^ from the xylem into the cell. This process prevents the excessive accumulation of toxic Na^+^ in the shoot and reproductive organs.

## SUPPLEMENTARY DATA

**Supplementary Fig. S1.** Salt hypersensitive phenotype of *sga* mutant grown *in vitro* at different salt concentrations.

**Supplementary Fig. S2.** *In vivo* phenotype of salt hypersensitive *sga1* and *sga2* mutants.

**Supplementary Fig. S3.** *In vitro* characterization of salt sensitive *sga* mutants.

**Supplementary Fig. S4.** The *sga1* mutant maintains similar leaf water loss to WT but increases the water contents and osmotic potentials in salt-treated leaves.

**Supplementary Fig. S5.** Spatial distribution of Na^+^ accumulation induced by salt stress in *sga1* mutant plants.

**Supplementary Fig. S6.** Expression levels of aquaporins (AQPs) genes in WT plants.

**Supplementary Fig. S7.** The salt hypersensitivity of adult grafted plants is only detected when *sga1-2* mutants are used as rootstocks.

**Supplementary Fig. S8.** The salt sensitivity is present in adult *sga1* mutant plants.

**Supplementary Table S1.** Oligonucleotides sequences used for the molecular cloning of the *sga1-2* mutations and generation of the CRISPR/Cas9 lines.

**Supplementary Table S2.** Sequences of primers for RT-qPCR gene expression analysis.

**Supplementary Table S3.** Phenotype inheritance in tomato *sga1* mutant under salt stress conditions.

**Supplementary Table S4.** Phenotype inheritance in tomato *sga2* mutant under salt stress conditions.

**Supplementary Table S5.** Co-segregation analysis between phenotype and a T-DNA insert with a functional *nptII* gene in tomato *sga1* and *sga2* mutants.

## ACKNOWLEDGEMENTS

Acknowledgement to the late Professor Bolarín MC for her invaluable contribution of vast experience in the conceptualization of this study and design of the experiments.

## AUTHORS CONTRIBUTION

IE, TB-L, YE and MJ-G performed genetic, molecular and physiological experiments and participated in the analysis and interpretation of experimental results. FAP, AA, BG-S, CC, JME-S, FB and TA collaborated in data analysis and experimental designs. IE, BP, FY-L, RL and VM contributed to conceptualization, directed the experiments and wrote the manuscript. All authors have critically reviewed the manuscript.

## DATA AVAILABILITY

All data can be found in the manuscript and in the supporting information.

## ABBREVIATIONS

AQPs: Aquaporins
DPI: Diphenylene Iodonium
DST: Days of Salt Treatment
FOX: Ferrous Ammonium Sulphate/Xylenol Orange
PIP: Plasma Membrane Intrinsic Proteins
ROS: Reactive Oxygen Species
TIP: Tonoplast Intrinsic Protein
WT: Wild Type

